# Tetrahydrocannabinolic Acid a (THCA-A) Reduces Adiposity and Prevents Metabolic Disease Caused by Diet-Induced Obesity

**DOI:** 10.1101/622035

**Authors:** Belén Palomares, Francisco Ruiz-Pino, Martin Garrido-Rodriguez, M. Eugenia Prados, Miguel A. Sánchez-Garrido, Inmaculada Velasco, María J. Vazquez, Xavier Nadal, Carlos Ferreiro-Vera, Rosario Morrugares, Giovanni Appendino, Gaetano Morello, Marco A Calzado, Manuel Tena-Sempere, Eduardo Muñoz

## Abstract

Cannabis has remarkable therapeutic potential, but its clinical use is limited by the psychotropic activity of Δ^9^-tetrahydrocannabinol (Δ^9^-THC). Surprisingly, the biological profile of the non-narcotic native precursor of Δ^9^-THC (Δ^9^-THC acid A, Δ^9^-THCA-A) is still largely unexplored. We present evidence that Δ^9^-THCA-A is a partial and selective PPARγ modulator, endowed with lower adipogenic activity than the full PPARγ agonist rosiglitazone (RGZ) and with an enhanced osteoblastogenic activity in human mesenchymal stem cells. Docking and *in vitro* functional assays indicated that Δ^9^-THCA-A binds to and activates PPARγ by acting at both the canonical and the alternative sites of the ligand-binding domain. Transcriptomic signatures at inguinal white adipose tissue (iWAT) from mice treated with Δ^9^-THCA-A confirmed its mode of action on PPARγ. Administration of Δ^9^-THCA-A in a mouse model of high fat diet (HFD)-induced obesity significantly reduced fat mass and body weight gain, markedly ameliorating glucose intolerance and insulin resistance, and largely preventing liver steatosis, adipogenesis and macrophage infiltration in fat tissues. Additionally, immunohistochemistry, transcriptomic, and plasma biomarker analyses showed that treatment with Δ^9^-THCA-A caused browning of iWAT and displayed potent anti-inflammatory actions in HFD mice. Altogether, our data validate the potential of Δ^9^-THCA-A as a low adipogenic PPARγ agonist, capable of substantially improving the symptoms of obesity-associated metabolic syndrome and inflammation. These findings suggest that Δ^9^-THCA-A, and perhaps non-decarboxylated *Cannabis sativa* extracts, are worth considering for addition to our inventory of cannabis medicines.

**SIGNIFICANCE STATEMENT:** The medicinal use of Cannabis is gaining momentum, despite the adverse psychotropic effects of Δ^9^-THC, the decarboxylation product of its naturally occurring and non-psychotropic precursor Δ^9^-THCA-A. We present evidence that Δ^9^-THCA-A is a partial ligand agonist of PPARγ with lower adipogenic activity compared to the full PPARγ agonist rosiglitazone (RGZ). Moreover, chronic administration of Δ^9^-THCA-A in a mouse model of high fat diet (HFD)-induced obesity significantly reduced body weight gain and fat mass, improved glucose intolerance and insulin resistance, and prevented liver steatosis and macrophage infiltration in fat tissues, additionally inducing white adipose tissue browning. Collectively, these observations qualify Δ^9^-THCA-A, a compound devoid of psychotropic effects, as an efficacious pharmacological agent to manage metabolic syndrome and obesity-associated inflammation.

**Highlights:**

- Δ^9^-THCA-A is a partial PPARγ ligand agonist with low adipogenic activity
- Δ^9^-THCA-A enhances osteoblastogenesis in bone marrow derived mesenchymal stem cells.
- Δ^9^-THCA-A reduces body weight gain, fat mass, and liver steatosis in HFD-fed mice
- Δ^9^-THCA-A improves glucose tolerance, insulin sensitivity, and insulin profiles *in vivo*
- Δ^9^-THCA-A induces browning of iWAT and has a potent anti-inflammatory activity

## INTRODUCTION

Preparations of *Cannabis sativa* have been used since the earliest written records of medical history, complementing the nutritional and technological uses of the plant and contributing to make it one of the first species cultivated by humans. Modern studies as well as anecdotal reports suggest the potential efficacy of cannabis extracts and cannabinoids in different medical conditions. Almost 150 cannabinoids have been isolated from different strains of Cannabis (1) with Δ^9^-tetrahydrocannabinol (Δ^9^-THC) and cannabidiol (CBD) being the best investigated class members from a medical standpoint. These neutral cannabinoids are produced and stored in the plant as acidic precursors (cannabinoid acids) (2). Decarboxylation requires heating but can take place also at room temperature upon prolonged storage of *Cannabis* (3). Remarkably, the acidic precursor of Δ^9^-THC (Δ^9^-THCA-A) is not psychotropic, and its binding to cannabinoid receptors is highly controversial and probably associated to contamination by its decarboxylation product (4). On the other hand, we recently showed that THCA-A, but not its decarboxylation product Δ^9^-THC, is a potent activator of the peroxisome proliferator-activated receptor-γ (PPARγ) endowed with remarkable neuroprotective activity (5). In agreement with these findings, the PPARγ activating properties of a Cannabis extract with a high titer of Δ^9^-THC was greatly reduced by decarboxylation (5). Given the key metabolic roles of PPARγ, these findings provided a rationale to investigate the possibility that Δ^9^-THCA-A based therapies could be beneficial for the management of metabolic disorders.

Obesity has reached pandemic proportions worldwide, with more than 650 million obese and nearly 2 billion people being overweighed (6). Obesity is a major risk factor for the development of multiple diseases, including prominently cardiovascular and metabolic disorders, such as type-2 diabetes and metabolic syndrome (MetS) (7, 8), but also respiratory, osteoarticular, cognitive, reproductive and oncologic pathologies (9–12). The basis for the multi-organ alterations seen in obesity remains unfolded (13), but the state of chronic, low-grade inflammation commonly associated to this condition is considered a major contributing factor (11). This is reflected by the increased concentrations of white adipose tissue-derived pro-inflammatory cytokines (TNF-α, IL-6, leptin) and the antiparallel decrease of anti-inflammatory signals, as adiponectin, commonly observed in obesity (12, 14, 15). Additional hormonal perturbations, e.g., of gastro-intestinal factors, such as ghrelin and GLP-1 (16), as well as the ectopic deposition of fat, mostly in the liver (i.e., steatosis), which defines a state of lipotoxicity (17), contribute also to the worsening of the metabolic profile of obese patients. In view of the conspicuous lack of effective therapies for the integral management of obesity, considerable attention is currently given to the identification of novel pharmacological agents globally targeting its deregulated substrates and other related pathways, as thermogenesis, to regain energy homeostasis (18–20).

In fact, while initial strategies to tackle obesity were mostly focused in the control of food intake, growing interest has been paid recently to elucidate the physiological roles and eventual therapeutic use of adaptive thermogenesis in the control of body weight and energy homeostasis (21, 22). Studies in this area have been directed not only to the brown adipose tissue (BAT), where canonical adaptive thermogenesis occurs in mammals, including humans (23), but also to the capacity of the white adipose tissue (WAT) to undergo, under certain conditions, browning, a process whereby adipocyte precursor cells acquire brown-like features, becoming beige or brite adipocytes (24). Brown and beige adipocytes are defined by a large set of mitochondria and high expression of uncoupling protein-1 (UCP-1), as major molecular switch to enhance heat dissipation at the expense of lower ATP synthesis at the respiratory chain (25). Thermogenic activity is finely regulated by the direct actions of a large set of circulating hormones, from adipokines to sex steroids and thyroid hormones (23, 26), as well as the adrenergic input from the sympathetic nervous system (27), therefore being amenable for pharmacological manipulation.

PPARγ is a nuclear receptor that plays key role in regulating a number of biological functions including lipid and glucose metabolism (28). PPARγ is also involved in the natural and adaptive immune response as well as in additional biological activities like the browning of white fat (29). PPARγ ligands include a wide array of natural and synthetic molecules, with glitazones, a class of compounds extensively for the management of type-2 diabetes, being the archetypal activators. Glitazones, like rosiglitazone (RGZ), are considered full PPARγ ligand agonists, and this profile is inextricably associated to both antidiabetic activity and undesirable side effects like weight gain, edema, and osteoporosis (30). Furthermore, liver injury, cancer, as well as an increased risk of heart failure have also been associated with the long-term use of glitazones (30). However, while the therapeutic potential of PPARγ ligand agonist remains high, interest has substantially shifted towards partial ligands, preeminent examples of which are natural products such as armofrutins and acidic cannabinoids (5, 31).

Depending on the mode of interaction and binding to the ligand-binding domain (LBD) of PPARγ (32, 33), modulators can induce graded responses, such as full- and partial agonism and antagonism. Thus, the recent identification of a functional alternative binding site for PPARγ ligands that does not fully overlap with the canonical glitazone binding site (34) has opened novel research avenues for the identification and/or development of second-generation PPARγ agonists.

## RESULTS

### Δ^9^-THCA-A is a selective and non-adipogenic PPARγ ligand agonist that induces iWAT browning through a PPARγ-dependent pathway

We have previously shown that Δ^9^-THCA-A is a PPARγ agonist at nanomolar concentrations (5). Herein, we have studied the selectivity of Δ^9^-THCA-A on different PPARs, showing that, when compared to the full ligand agonist rosiglitazone (RGZ), Δ^9^-THCA-A is only a partial ligand activator for PPARγ, devoid of PPARα and PPARδ transcriptional activities (**Fig. 1A**). Dephosphorylation of PPARγ at Ser273 is essential for acquiring insulin-sensitizing activity (35), and we found that, in 3T3-L1 adipocytes, Δ^9^-THCA-A, as well as RGZ, could inhibit the TNFα-induced Ser273 phosphorylation of PPARγ (**Fig. 1B**).

**Figure 1.**
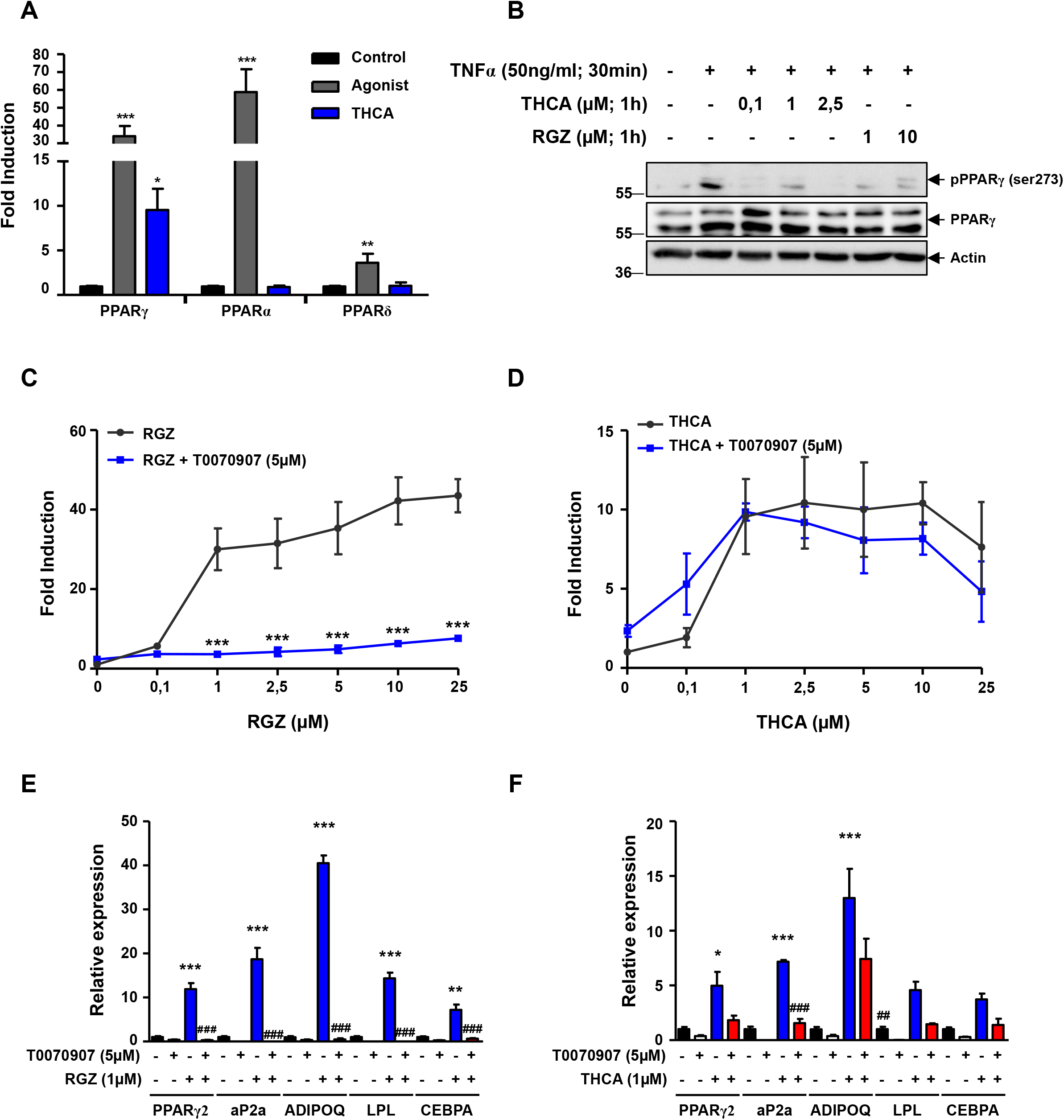
Characterization of Δ^9^-THCA-A as a selective PPARγ agonist. **(A)** Receptor-specific transactivation by Δ^9^-THCA-A. HEK-293T cells were co-transfected with the plasmids encoding nuclear receptors (GAL4-PPARγ, GAL4-PPARα and GAL4-PPARδ) and their cognate luciferase reporter (GAL4-luc). After transfection, cells were treated with Δ^9^-THCA-A (1 μM) and receptor-specific agonists for 6 hours. Control (black bars), VCE-004.8 (blue bars) and specific ligands for each receptor (grey bars): RGZ (1 μM) for PPARγ, WY14643 (5 μM) for PPARα, and GW0742 (5 μM) for PPARδ. Results are expressed as the fold induction ± SD (n = 3) relative to untreated control. **(B)** Representative Western blot images of PPARγ phosphorylation at Ser273 in 3T3L1 adipocytes pre-treated with Δ^9^-THCA-A and RGZ for 30 min, followed by treatment with TNF-α for 30min (n = 3). **(C, D)** HEK-293T cells were co-transfected with GAL4-PPARγ and GAL4-luc, pre-treated with T0070907 for 15 min and then stimulated with increasing concentrations of either RGZ or Δ^9^-THCA-A for 6 hours and assayed for luciferase activity. Results are expressed as the fold induction ± SD relative to RGZ **(C)** or Δ^9^-THCA-A **(D)** (n = 4). **(E, F)** Adipocyte differentiation of MSCs in adipogenic medium with RGZ **(E)** or Δ^9^-THCA-A **(F)** in the presence and the absence of T0070907 for 7 days. mRNA levels of adipogenic markers were analysed by qPCR. Results represent the mean ± S.D (n = 3). *P < 0.05, **P < 0.01 and ***P < 0.001 agonist ligands or Δ^9^-THCA-A treatment vs. control; RGZ + T0090709 vs. RGZ; RGZ or Δ^9^-THCA-A treatment vs. control; ^##^P < 0.01 and ^###^P < 0.001 RGZ + T0090709 vs RGZ or Δ^9^-THCA-A + T0090709 vs Δ^9^-THCA-A. (ANOVA followed by Tukey’s test or unpaired two-tailed Student’s t-test).

PPARγ has a large and dynamic LBD, whose diversity of ligand accommodation in its two sub-pockets is associated to distinct biological activities (32). Based on the structure of different PPARγ-ligand complex, it has been proposed that full agonists bind to both sub-pockets, establishing hydrogen bonds with residues Tyr473 (H12) on AF-2 (also called orthosteric or canonical binding site) and Ser342 (S1/S2) on β-sheet sub-pocket (also called alternative binding site), whereas partial agonists can preferentially bind to the alternative site (34). Docking analysis suggested that Δ^9^-THCA-A binds both the canonical (Ser289) and the alternative binding sites (L340 and Ser342) in LBD (**Fig. S1**). The functionality of both LBD sites in the response to Δ^9^-THCA-A was therefore investigated. To this purpose, T0070907, a PPARγ antagonist that binds irreversibly to the canonical binding site, was used. First, luciferase reporter assays were used to study the participation of the canonical and alternative binding sites in the response to Δ^9^-THCA-A in comparison with RGZ. As expected, preincubation with T0070907 effectively blocked RGZ-induced PPARγ transactivation (**Fig. 1C**). Conversely, T0070907 could not block Δ^9^-THCA-A-induced PPARγ transcriptional activity (**Fig. 1D**). In addition, T0070907 almost fully suppressed the expression of PPARγ-dependent genes induced by RGZ in human mesenchymal stem cells (MSCs) differentiated to adipocytes (**Fig. 1E**). Conversely, T0070907 did not fully prevent the effect of Δ^9^-THCA-A on these genes (**Fig. 1F**). By using time-resolved fluorescence resonance energy transfer (TR-FRET) co-regulator interaction assays, we found that RGZ increases the binding of peptides derived from TRAP220 and PGC-1α, but decreases the binding of peptides derived from NCoR and SMRT. In contrast, Δ^9^-THCA-A has not significant effect on the recruitment TRAP220 to PPARγ and showed only a moderate effect on the recruitment of PGC-1α. Indeed, Δ^9^-THCA-A had a weak effect to displace corepressors from the binding to PPARγ (**Fig. S2**).

To further investigate the role of the PPARγ canonical binding site *in vivo*, mice were treated with Δ^9^-THCA-A in the presence or not of T0070907, performing transcriptomic analysis in inguinal white adipose tissue (iWAT), a major target organ for PPARγ agonists. The comparative differential expression analysis of both conditions versus control mice resulted in a total of 3719 genes with an adjusted P ≤ 0.01 and an absolute fold change ≥ 2 (**Fig. 2A**). From them, 72 genes belonging to the PPARγ signaling and thermogenesis pathways were upregulated in response to Δ^9^-THCA-A (**Fig. 2A-B**). Within this group of genes, the co-treatment with T0070907 prevented the upregulation of 46 genes induced by Δ^9^-THCA-A and 26 genes were unaffected (**Fig. 2C**). In contrast, genes included in the NF-κB signaling pathway were not affected by Δ^9^-THCA-A treatment, but significantly increased their expression in animals co-treated with T0070907 (**Fig. 2A-B**). Uncoupling protein 1 (UCP-1) gene, a key marker of the iWAT browning process, was found among the top 25 genes upregulated by Δ^9^-THCA-A in a T0070907-sensitive manner (**Fig. 2D**). Moreover, western blotting and immunohistochemistry showed that Δ^9^-THCA-A induced the expression of UCP-1 protein in iWAT (**Fig. 2E**). In line with such putative thermogenic activation, Δ^9^-THCA-A treatment caused a suppression of body weight that was independent of food intake changes but blocked by co-administration of T0070907 (**Fig. S3**).

**Figure 2.**
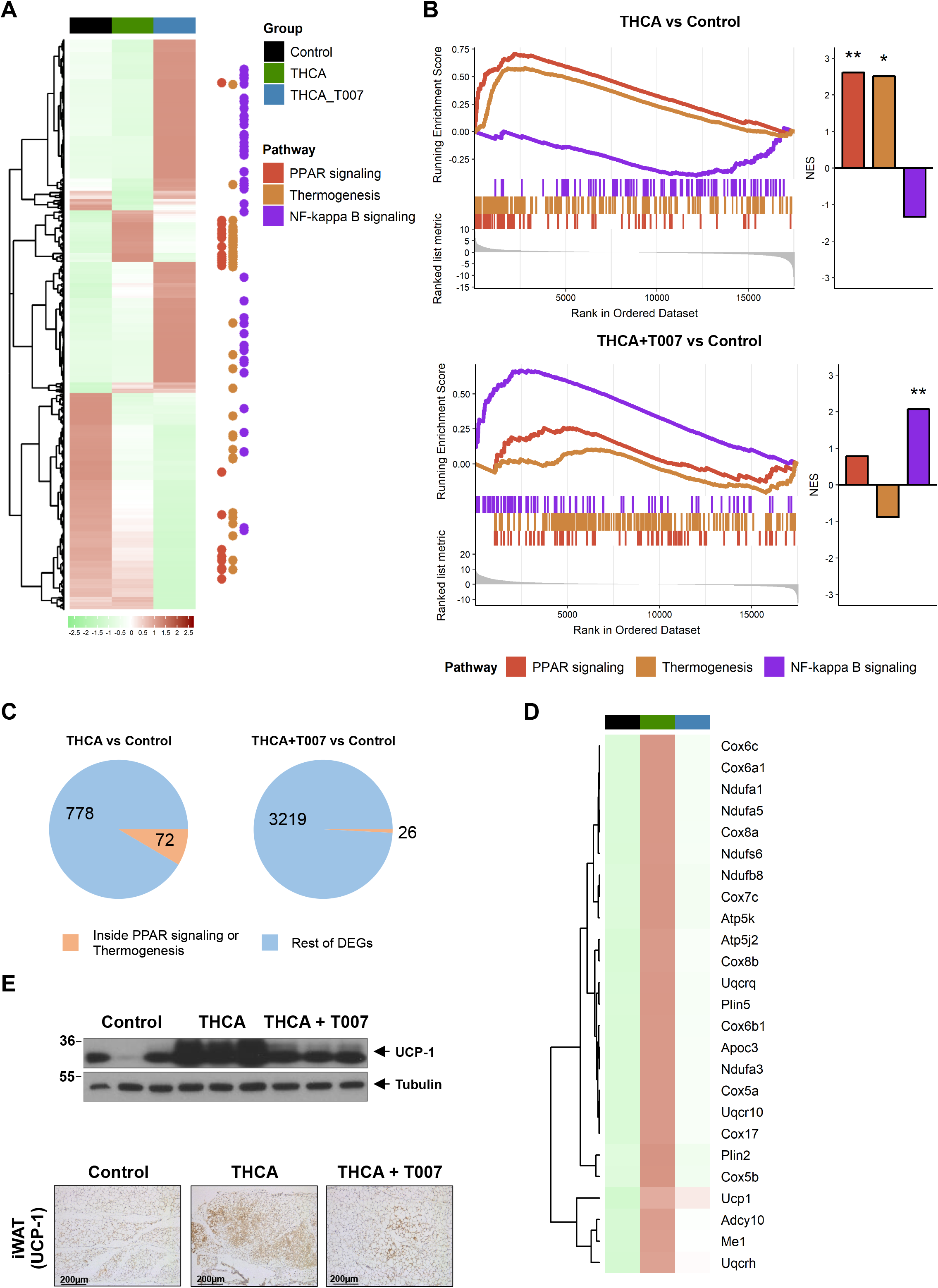
Transcriptomic analysis of the PPARγ dependent activity of Δ^9^-THCA-A in iWAT. **(A)** Heatmap of all the differentially expressed genes (absolute fold change ≥ 2 and an adjusted P value ≤ 0.01) in Δ^9^-THCA-A versus control or Δ^9^-THCA-A+T0070907 versus control comparisons. The color represents the scaled mean of log transformed expression. The column annotations indicate the sample group and the points at the right side highlight the position of genes belonging to the KEGG pathways of interest. **(B)** Gene set enrichment analysis results for the KEGG pathways of interest. The left side enrich plots indicate the position of the genes belonging to each pathway in the pre-ranked list per comparison. The right-side bar plots represent the normalized enrichment score (NES) and significance of the GSEA result *P ≤ 0.05; **P ≤ 0.01; ***P ≤ 0.001. **(C)** Pie charts indicating the number and proportion of differentially expressed genes included in the KEGG pathways of interest for each comparison. **(D)** Heatmap of the top 25 genes induced by Δ^9^-THCA-A inside the PPAR signaling or thermogenesis pathways that are not differentially expressed in the Δ^9^-THCA-A+T0070907 vs control comparison. **(E)** Representative Western blot images of UCP-1 protein expression and immuno-histochemistry with anti-UCP-1 antibodies in iWAT tissue (original magnification × 10, scale bar: 200 μm) (n = 3).

In our analysis, we also found 317 genes that were modified by Δ^9^-THCA-A in a way insensitive to T0070907 (**Fig. S4A**). To functionally evaluate this cluster of genes, we performed an over-representation analysis using the KEGG pathways and Gene Ontology annotations after sorting them out into up- and down-regulated ones (**Fig. S4B-C**). Interestingly, a set of upregulated genes associated to fatty acid metabolism was found. Conversely, genes belonging to cGMP-PKG, to calcium signaling pathways, and to muscle tissue development were downregulated by Δ^9^-THCA-A administration. Taken together, our results showed that the bioactivity of Δ^9^-THCA-A was mediated by the PPARγ canonical pathway as well as by pathways mediated by the alternative binding site of this nuclear receptor.

PPARγ partial ligand agonists are thought to be less adipogenic than full ligand agonists and are also expected to have fewer negative effects on bone metabolism. Therefore, we studied the ability of Δ^9^-THCA-A to influence MSCs differentiation into adipocytes and osteoblasts. MSCs were cultured in adipogenic medium (AM) for either 14 days or 21 days to evaluate the mRNA expression of adipogenic markers and to detect lipid droplets. **Fig. 3A-B** show that MSC treated with Δ^9^-THCA-A contained fewer and smaller lipid droplets compared to RGZ treatment. Moreover, Δ^9^-THCA-A induced lower expression of the adipogenic differentiation markers PPARγ, aP2a, ADIPOQ, LPL and CEBPA, as compared to cells treated with RGZ (**Fig. 3C**). Remarkably, we found that, in MSC differentiated in an osteoblastogenic medium, Δ^9^-THCA-A enhanced osteoblast mineralization as well as the expression of the osteogenic differentiation markers Runx2, SP7, IBS and ALP, (**Fig. 3D-F**). These data indicate that Δ^9^-THCA-A qualifies as a partial PPARγ ligand significantly less adipogenic than RZG and with an enhanced osteoblasts differentiation capacity.

**Figure 3.**
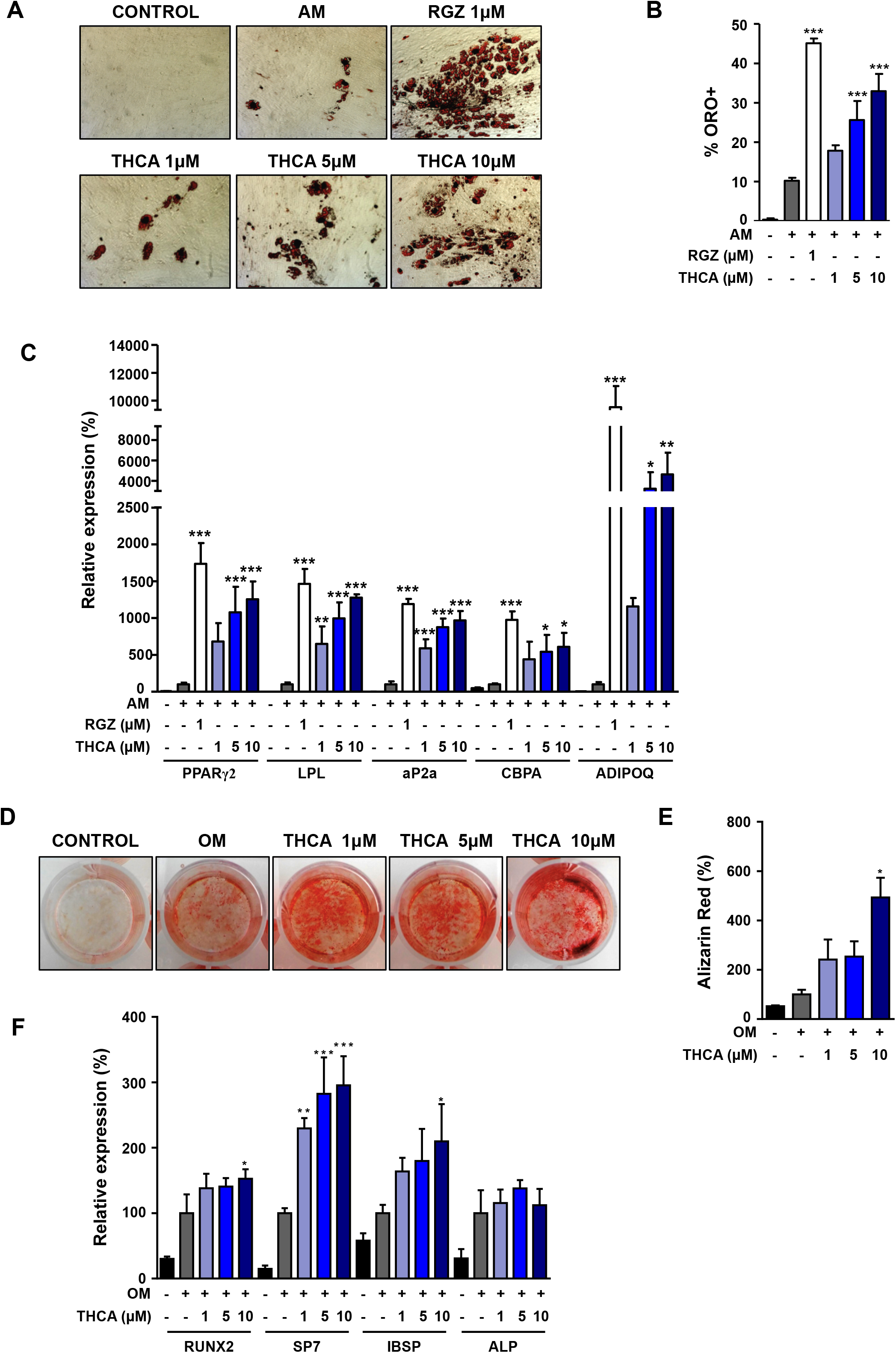
Effect of Δ^9^-THCA-A on MSCs adipogenic and osteoblastogenic differentiation. MSCs were cultured under adipogenic medium (AM) in the presence of RGZ or Δ^9^-THCA-A. **(A)** Representative images of the cells stained with Oil Red O assayed by light microscopy (×10) after 21 days of differentiation. **(B)** Quantification of the stained lipid droplets were performed measuring absorbance at 540nm. **(C)** mRNA levels of adipogenic markers were analysed by qPCR after 14 days of differentiation. MSCs were cultured under osteoblastogenic medium (OM) in the presence of Δ^9^-THCA-A. **(D)** Mineralization detected by Alizarin red staining was assessed by gross appearance after 21 days of differentiation. **(E)** Quantification of the eluted Alizarin Red stain measuring absorbance at 405nm. **(F)** Gene expression of osteoblastogenic markers analysed by qPCR. Results represent the mean ± S.D (n = 3). *P < 0.05, **p < 0.01 and ***P < 0.001 RGZ or Δ^9^-THCA-A vs. AM; Δ^9^-THCA-A vs. OM. (ANOVA followed by Tukey’s test).

### Δ^9^-THCA-A treatment ameliorates HFD-induced metabolic perturbations and iWAT inflammation

In addition, we analyzed the effects of chronic treatment for 3-wks with an effective dose of Δ^9^-THCA-A (20 mg/kg BW/day) on various metabolic and hormonal parameters in a mouse model of HFD-induced obesity. Feeding a HFD for 12-wks resulted in a significant increase in body weight over CD controls (BW; **Fig. 4A**), together with enhanced fat mass (35.89 ± 2.63 g vs. 16.19 ± 1.45 g in CD; P>0.001) and adiposity index, calculated as ratio between fat mass and fat + lean mass (36.43 ± 2.62 vs. 16.56 ± 1.49 % in CD; P=0.001). Repeated administration of Δ^9^-THCA-A caused a marked suppression of BW gain in HFD mice (**Fig. 4B**), as observed, albeit with lesser amplitude, in CD mice. The increased adiposity caused by HFD was similarly restored by chronic Δ^9^-THCA-A treatment, which caused also a significant suppression of the adiposity index in control lean mice (**Fig. 4C**).

**Figure 4.**
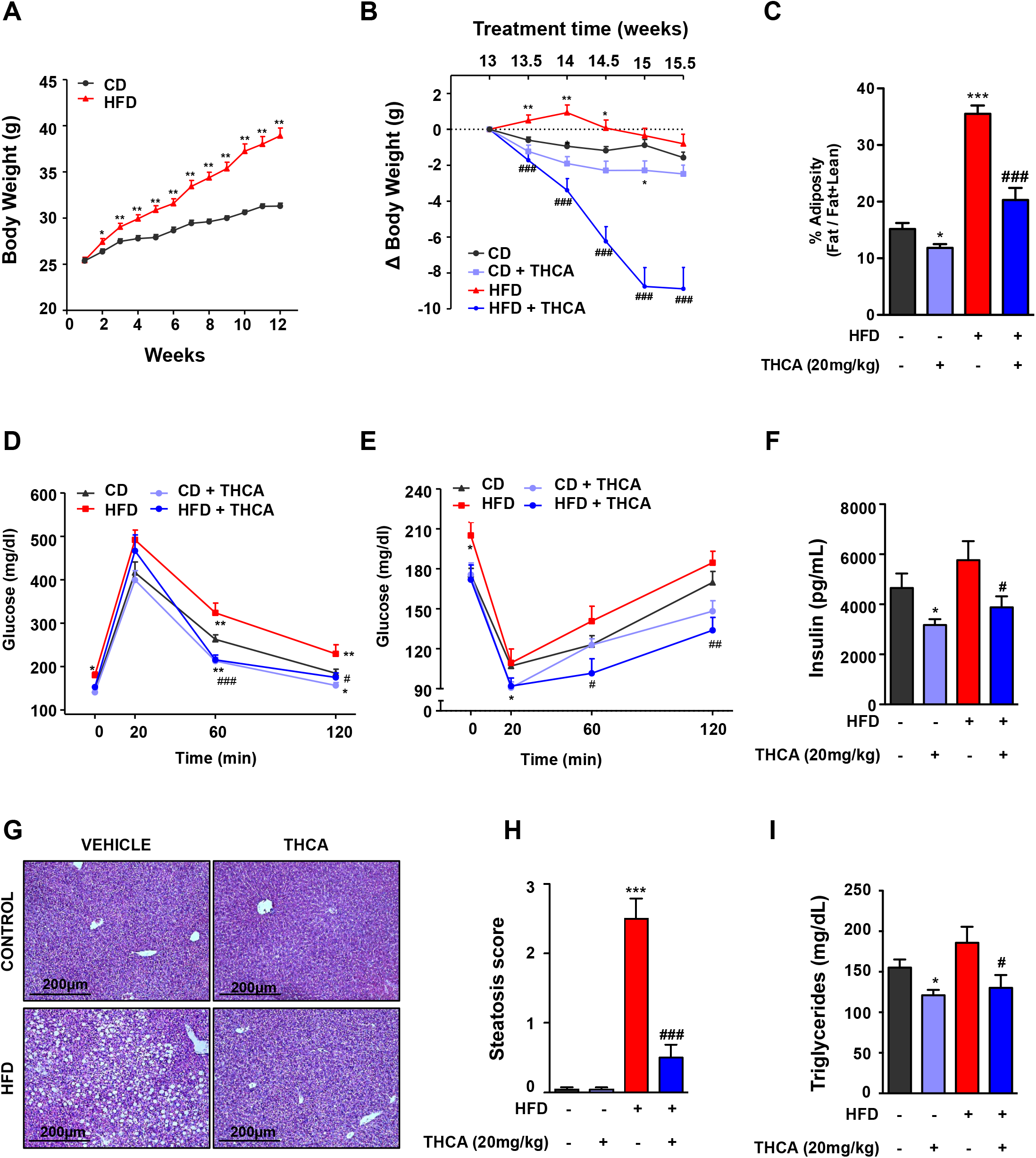
Effect of administration of Δ^9^-THCA-A on metabolic and hormonal parameters in a mouse model of HFD-induced obesity. **(A)** Body weight (BW) evolution of adult male mice fed for 12-weeks with high fat diet (HFD) or the corresponding control diet (CD). **(B)** BW change in HFD and CD mice treated for three weeks with Δ^9^-THCA-A or vehicle; values are referenced to BW at the beginning of treatment (taken as 0). **(C)** Percentage of adiposity, at the end of treatments in the four experimental groups. **(D-E)** Glucose and Insulin tolerance tests in CD and HFD mice treated with Δ^9^-THCA-A or vehicle for three weeks. **(F)** Basal insulin levels at the end of the three-week treatment period are shown for the four experimental groups. **(G)** Liver sections with hematoxylin and eosin (H&E) staining (original magnification x10, scale bar: 200 μm). **(H)** Steatosis scores (n = 6 mice per group) and **(I)** plasma levels of triglycerides. Values correspond to means ± SEM of at least 8 mice per group. \**P*<0.05, \*\**P*<0.01, \*\*\**P*<0.001 Δ^9^-THCA-A-treated mice or HFD mice vs. control (CD) mice; ^#^*P*<0.05, ^##^*P*<0.01, ^###^*P*<0.001 Δ^9^-THCA-A-treated HFD mice vs. HFD mice treated with vehicle (ANOVA followed by Tukey’s test or unpaired two-tailed Student’s t-test).

Administration of Δ^9^-THCA-A for 3-wks improved also glucose homeostasis in HFD-induced obese mice. Thus, while HFD exposure for 15-wks evoked an elevation of basal glucose levels, worsened glucose tolerance after glucose bolus injection (**Fig. 4D**) and reduced insulin sensitivity (**Fig. 4E**), treatment of HFD mice with Δ^9^-THCA-A for 3-wks resulted in lowering of basal glycemia, and markedly improved glucose profiles, both in glucose and insulin tolerance tests, which displayed better profiles than those of CD mice without pharmacological intervention. Moreover, positive effects of Δ^9^-THCA-A in terms of glucose tolerance and insulin sensitivity were also detected in lean control mice (**Fig. 4D-E**). This was associated to a significant lowering of basal insulin levels after Δ^9^-THCA-A administration to CD and HFD mice (**Fig. 4F**), possibly reflecting a state of enhanced insulin sensitivity. In the same line, Δ^9^-THCA-A treatment of obese mice largely prevented liver fat infiltration caused by HFD and markedly reduced the steatosis score (**Fig. 4G-H**). In good agreement with these observations, Δ^9^-THCA-A significantly decreased serum triglyceride levels both in obese and lean mice (**Fig. 4I**).

The iWAT transcriptomic profile in control and HFD mice, untreated or treated with Δ^9^-THCA-A, was next investigated. A differential expression analysis revealed a total of 1387 genes overcoming the cutoff of an adjusted P ≤ 0.01 and an absolute fold change ≥ 2 in any of the two comparisons (**Fig. 5A**). Among them, KEGG pathway analyses revealed that genes involved in NF-κB signaling and cytokine-cytokine receptor were upregulated in HFD mice, which matches the inflammatory phenotype in iWAT that accompanies the metabolic syndrome (36). Additionally, genes belonging to the insulin receptor signaling pathway showed a lowered expression in HFD mice, in agreement with the hallmark state of insulin resistance of obesity. Interestingly, Δ^9^-THCA-A treatment in HFD mice reduced the expression of genes belonging to the inflammatory pathways, partially recovering the expression of those linked to the insulin signaling process (**Fig. 5A-B**). In total, DEGs analysis revealed that 1014 genes including those related to NF-κB signaling and cytokine-cytokine receptor (70 genes) were modified in HFD mice and normalized in Δ^9^-THCA-A-treated HFD mice (**Fig. 5C**). Finally, to confirm the anti-inflammatory profile of Δ^9^-THCA-A, we analyzed by qPCR assays the top 25 upregulated inflammatory genes in the iWAT of HFD mice (**Fig. 5D**), confirming the increased expression of key components of this gene set, including TNFα, ICAM-1, CD4, CXCL-16, CCK22, CXCR5 and CXCR2 (**Fig. 5E**).

**Figure 5.**
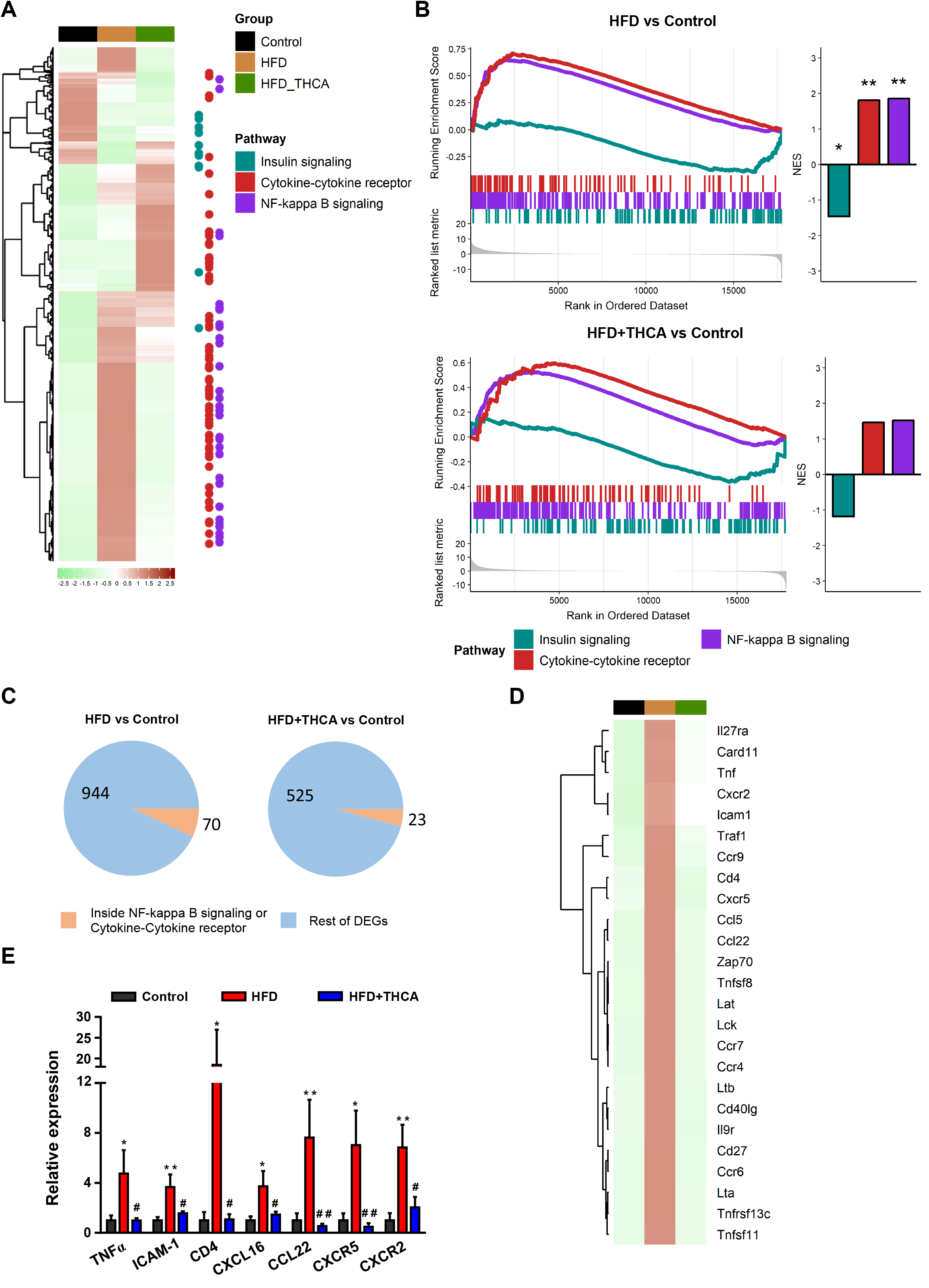
Transcriptomic analysis of Δ^9^-THCA-A effects in the iWAT of HFD mice. **(A)** Heatmap of all the differentially expressed genes (absolute fold change ≥ 2 and an adjusted P value ≤ 0.01) in HFD versus control or HFD+Δ^9^-THCA-A versus control comparisons. The color represents the scaled mean of log transformed expression. The column annotations indicate the sample group and the points at the right side highlight the position of genes belonging to the KEGG pathways of interest. **(B)** Gene set enrichment analysis results for the KEGG pathways of interest. The left side enrich plots indicate the position of the genes belonging to each pathway in the pre-ranked list per comparison. The right-side bar plots represent the normalized enrichment score (NES) and significance of the GSEA result *P ≤ 0.05; **P ≤ 0.01; ***P ≤ 0.001. **(C)** Pie charts indicating the number and proportion of differentially expressed genes included in the KEGG pathways of interest for each comparison. **(D)** Heatmap of the top 25 genes induced by HFD inside the NF-κB or cytokine-cytokine receptor pathways that are not differentially expressed in the HFD+Δ^9^-THCA-A vs control comparison. **(E)** Gene expression of pro-inflammatory genes were measured by qPCR. Results are presented as mean ± SEM of at least 5 mice per group. *P<0.05, **P<0.01 HFD mice vs. control mice; #P<0.05, ##P<0.01 Δ9-THCA-A-treated HFD mice vs. HFD mice treated with vehicle (ANOVA followed by Tukey’s test).

In keeping with their obese phenotype, HFD mice also displayed features of adipose tissue enlargement and inflammation. Thus, a significant increase in adipocyte volume was observed, accompanied by signs of macrophage infiltration of white adipose tissue (WAT), assessed by F4/80 staining of clusters of macrophages surrounding dead adipocytes, in the so-called crown-like structures (CLS) (**Fig. 6A-B**). Notably, Δ^9^-THCA-A administration for 3-wks fully prevented adipocyte enlargement and the appearance of CLS in the inguinal WAT of HFD-induced obese mice. Moreover, a substantial upsurge in the levels of the thermogenic protein, UCP-1, in the iWAT of obese mice was also observed (**Fig. 6A-C**).

**Figure 6.**
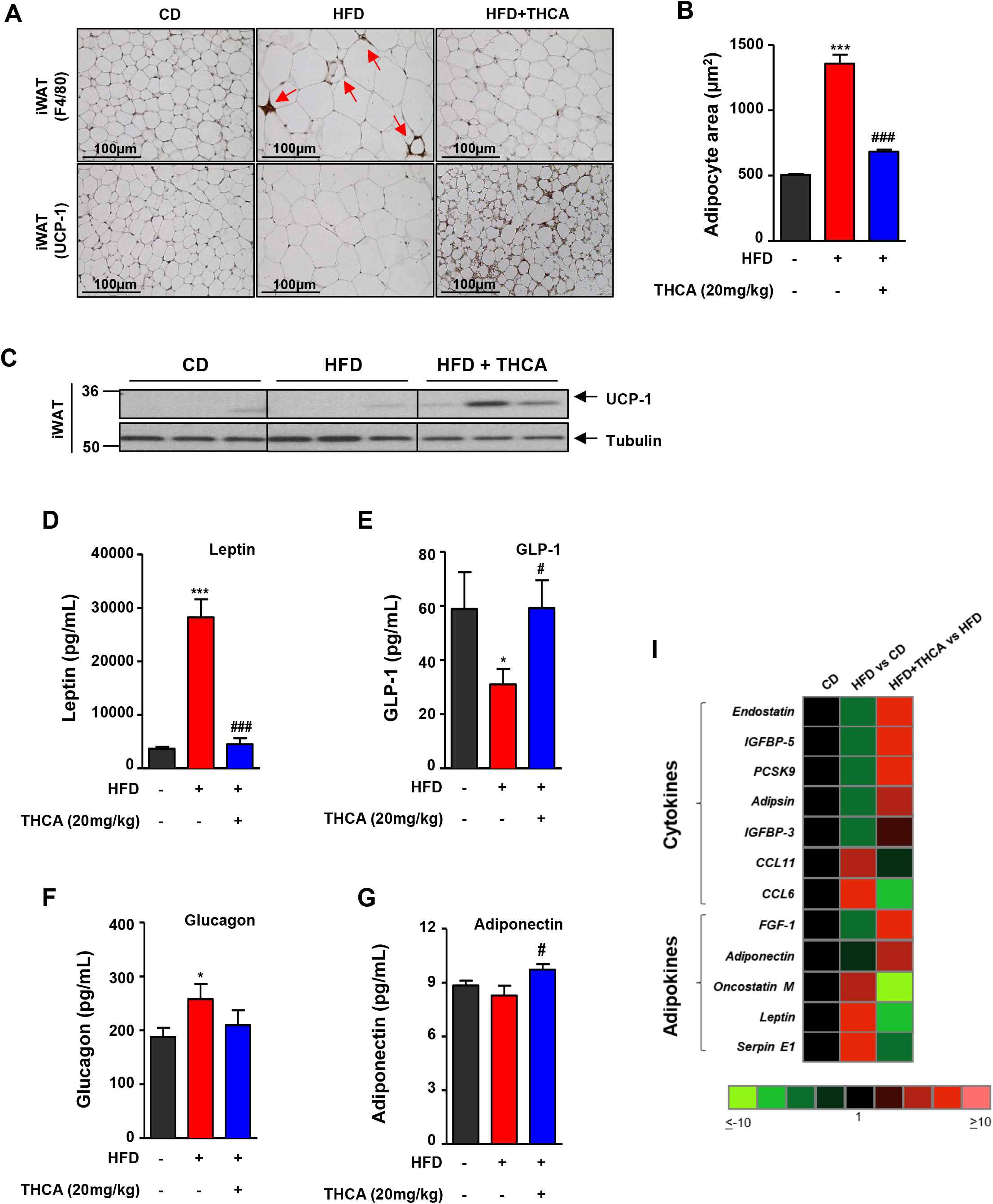
Effects of Δ^9^-THCA-A on iWAT browning, adiposity and circulating factors, with key roles in metabolic homeostasis in CD and HFD animals. **(A)** Crown Like Structures (CLS) and browning in iWAT. Representative immunohistochemical detection of anti-F4/80 and anti-UCP-1 antibodies (original magnification x20, scale bar: 100 μm), **(B)** Quantification of adipocyte area (n = 6 animals per group), **(C)** UCP-1 protein levels determined by western blotting in iWAT tissue (n=3). Hormonal markers linked to energy and metabolic homeostasis assayed: **(D)** Leptin, (E) GLP-1, **(F)** Glucagon and **(G)** Adiponectin. **(I)** Heatmap showing the plasma profile of cytokines and adipokines. Values correspond to means ± SEM of at least 8 mice per group. \**P*<0.05, \*\*\**P*<0.001 HFD mice vs. control (CD) mice; ^#^*P*<0.05, ^###^*P*<0.001 Δ^9^-THCA-A-treated HFD mice vs. HFD mice treated with vehicle (ANOVA followed by Tukey’s test or unpaired two-tailed Student’s t-test).

Finally, changes in the circulating levels of key metabolic hormones were assessed in HFD-induced obese mice, treated or not with Δ^9^-THCA-A. In line with their obese phenotype, HFD mice showed increased leptin and glucagon levels, and decreased GLP-1 concentrations vs. lean CD animals. Treatment with Δ^9^-THCA-A for 3-wks fully normalized leptin, GLP-1 and glucagon levels, and evoked a significant increase in adiponectin concentrations (**Fig. 6D-G**). In the same vein, proteome profiler arrays targeting a comprehensive set of circulating adipokines and cytokines confirmed the increase of the levels of leptin, together with other adipose born-factors, such as Oncostatin M and Serpin E1, in HFD mice, which were decreased by Δ^9^-THCA-A, while adiponectin concentrations were increased. Likewise, a set of cytokines including Endostatin, IGFBp-5, PCSK9, Adipsin and IGFBP-3 were suppressed by HFD but increased by Δ^9^-THCA-A in HFD mice, while an opposite pattern was observed for CCL11 and CCL6 (**Fig. 6I**).

## DISCUSSION

Because of its escalating prevalence and the enormous health and economic burden associated to its co-morbidities, such as type 2 diabetes, obesity has become a major societal problem, now recognized as the major preventable risk factor for all-cause mortality (6, 37), with a nearly 50% more life-years lost compared to other major avertable factors, like smoking. Glitazones, such as RGZ, are synthetic full PPARγ agonists that have been marketed since the early 2000s for the treatment of type 2 diabetes. However, the use of glitazones in diabetic patients has dropped significantly over the past years due to a series of adverse side effects that include bone loss and osteoporosis, as well as fluid retention. Bone loss could be bound to the effect of glitazones on the lineage commitment of mesenchymal stem cells towards osteoblasts and adipocytes, and it has been shown that RGZ suppresses osteoblastogenesis and induces adipocyte differentiation (38). The development of more balanced partial PPARγ agonists, devoid of the side effects showed by the currently marketed PPARγ full agonists, is considered a major challenge for drug discovery (39). We provide evidence that Δ^9^-THCA-A is a partial PPARγ agonist, lacking the adverse psychotropic effects of Δ^9^-THC, with a lower adipogenic capacity than glitazones, and endowed also with osteoblastogenesis-enhancing properties. In *in vivo* animal experiments, Δ^9^-THCA-A could successfully prevent adiposity and reversed the metabolic and inflammatory complications associated to diet-induced obesity.

Docking and functional analysis revealed that Δ^9^-THCA-A may exert its PPARγ activity by acting at both the canonical and the alternative binding sites of the PPARγ LBD. Δ^9^-THCA-A binds to the alternative site by interacting with Ser342 at the β-sheet (Ω-loop) through its carboxylate group. This may explain the observation that Δ^9^-THCA-A is at least 20-fold more potent than Δ^9^-THC to activate PPARγ (5). In addition, dietary long- and medium-chain fatty acids, as well as small molecules, also bind to the same amino acid in the alternative site, highlighting the relevance of this site for a wide class of synthetic and natural compounds including acidic cannabinoids (40, 41). Our data also suggest that Δ^9^-THCA-A may stabilize the Ω-loop region and inhibit phosphorylation of PPARγ Ser273, thus inducing anti-diabetic activity. However, Δ^9^-THCA-A does not affect PPARγ co-regulators interaction, and we cannot discard that Δ^9^-THCA-A could also act as a neutral ligand at the orthosteric site, since most of its activities are inhibited by T0070907. Thus, Δ^9^-THCA-A could compete with RGZ in PPARγ-dependent transcriptional assays (5), and TR-FRET assays also showed that Δ^9^-THCA-A partially prevented the recruitment of TRAP220 and SMRT induced by RGZ. In this model, Δ^9^-THCA-A could push endogenous fatty acids out of the orthosteric pocket toward the alternative binding site (40). Nevertheless, it has been shown that others partial agonists such as INT131 and BVT-13 do not induce significant binding or displacement of PPARγ co-regulators (32, 41, 42). Interestingly, SR2595, an inverse PPARγ agonist that blocks phosphorylation of Ser273, could promote osteoblastogenesis in cultured hMSCs (43). In summary, our *in vitro* and *in vivo* data strongly suggest that the binding of Δ^9^-THCA-A to the PPARγ orthosteric site accounts for most of the Δ^9^-THCA-A effects mediated by this nuclear receptor, but signaling through the alternative site could also be of biological relevance. Thus, in lean mice, 317 genes modified by Δ^9^-THCA-A but unaffected by T0070907 were found. Among them, upregulation of genes related with the fatty acid metabolism and downregulation of genes belonging to cGMP-PKG, calcium signaling pathways and muscle tissue development, were identified. Whether this pathway is linked or not to the PPARγ alternative site or to other pathways, such as CB_1_ signaling, remains to be investigated.

There is growing evidence of a link between obesity and inflammation, and Δ^9^-THCA-A showed a very potent anti-inflammatory profile in HFD mice. Thus, activation of innate and adaptive immune response has been observed in the fat tissue in obese individuals (44–46), and we identified 70 upregulated genes linked to NF-κB and cytokine-cytokine-receptor signaling, whose expression was largely prevented by treatment with Δ^9^-THCA-A. Within them, T- and B cell markers, as well as macrophage-derived cytokines and chemokines, were identified, showing that both the innate and the adaptive immune responses account for the inflammatory status in the fat tissue of HFD mice. There is strong evidence that PPARγ inhibits NF-κB activation through several mechanisms, repressing NF-κB-mediated transcription of proinflammatory cytokines in immune and non-immune cells (47). Thus, Δ^9^-THCA-A could dampen inflammation mainly through the PPARγ pathway, and this phytocannabinoid deserves consideration for the management of other chronic or acute inflammatory diseases.

The efficacy of Δ^9^-THCA-A to alleviate a wide spectrum of metabolic and hormonal derangements linked to diet-induced obesity was evaluated in a validated preclinical model of metabolic syndrome, namely, 12-wk exposure to HFD (48). This model fully recapitulates the cardinal manifestations of obesity and its major complications, including an increase of body weight, fat mass, adipocyte area and adiposity index, accompanied by perturbed glucose homeostasis and insulin resistance, as well as enhanced liver lipid deposition and steatosis. Notably, a regimen of 3-wk treatment with an effective daily dose of Δ^9^-THCA-A was sufficient to reverse such adverse metabolic profile, causing a marked suppression of body weight gain and adiposity, significantly lowering basal glucose and insulin levels, and fully preventing HFD-induced glucose intolerance and insulin resistance. Moreover, circulating triglyceride levels were reduced and steatosis scores were substantially improved by 3-wk Δ^9^-THCA-A treatment in HFD mice, substantially reversing the metabolic phenotype caused by the high fat content diet. Likewise, the major hormonal and cytokine alterations caused by HFD were reversed by Δ^9^-THCA-A, with a significant decrease in circulating leptin and glucagon, and a significant increase in serum GLP-1 and adiponectin levels. Such a switch towards an anti-inflammatory state was also detected by our adipo-/cytokine arrays, which confirmed the reversal of most of the pro-inflammatory humoral alterations caused by HFD. Interestingly, the beneficial effects of Δ^9^-THCA-A were not only observed in HFD conditions, but also, albeit to a lesser extent, in lean animals fed a control diet, in which 3-wk treatment with Δ^9^-THCA-A was capable to reduce body weight and adiposity, as well as basal glucose, insulin and triglyceride levels, together with a significant enhancement of glucose tolerance and insulin sensitivity. All these features define an optimal pharmacological profile of Δ^9^-THCA-A for globally handling the metabolic syndrome linked to obesity.

The beneficial effects of Δ^9^-THCA-A on body weight and metabolic status are somewhat at odds with the reported orexigenic and lipogenic activity of cannabinoids (49, 50). In our studies, Δ^9^-THCA-A failed to cause any detectable change in food intake, supporting a distinct pharmacological profile compared to Δ^9^-THC, whose feeding-promoting effects underlie its use in the management of cachexia (51). The lack of orexigenic effect together with the substantial body weight loss induced by Δ^9^-THCA-A suggests a potential use of this compound to induce adaptive thermogenesis. Indeed, chronic treatment with Δ^9^-THCA-A caused the upregulation of a large set of genes belonging of the thermogenesis pathway, with a prominent increase in the UCP-1 content in inguinal WAT that was largely prevented by cotreatment with the PPARγ inhibitor, T0070907. UCP-1 induction was also detected in the WAT of HFD mice. Altogether, these findings suggest that, acting at least partially via PPARγ, Δ^9^-THCA-A causes browning of the white adipose tissue, and this activity could mechanistically underlie the beneficial effects of this carboxylated cannabinoid on energy and metabolic homeostasis. These features add to our current efforts to identify pharmacological agents capable to activate thermogenesis without undesired side effects.

The global challenge posed by obesity and the inherent difficulties to handle its multi-factorial pathophysiology have fueled the search for novel therapeutic agents endowed with multi-target activity and capable to improve the different metabolic alterations associated to overweight. Recent efforts in this area include the development of novel hormonal multi-agonists based on peptide chimeras, or on conjugates fusing peptides and small molecules, capable to jointly target the various signaling pathways involved in the pathogenesis of the major complications of obesity (20). While this approach holds the promise to improve obesity treatments, it is not devoid of potential side effects and the pharmacodynamic limitations linked to the integration of different hormonally-active moieties with, in some cases, opposite biological activities. In the context of a poly-pharmacological approach to obesity and metabolic syndrome, the phytocannabinoid, Δ^9^-THCA-A, deserves further studies, since it is endowed with the beneficial effects of PPARγ agonists but devoid of their adipogenic activity and of the adverse psychotropic and orexigenic effects of narcotic cannabinoids. The effects of the administration of Δ^9^-THCA-A in a preclinical model of diet-induced obesity equal, if not outperform, the results reported for promising poly-agonist therapies recently advocated for obesity treatment (18–20), suggesting that this non-psychotropic phytocannabinoid, as well as non-decarboxylated *Cannabis sativa* extracts, are worth of consideration for the management of obesity and metabolic disease.

## Methods

### Δ^9^-THCA-A isolation

Δ^9^-THCA-A was purified at >95% from the proprietary Cannabis variety Moniek (CPVO/20160114) using a Counter Current Chromatography (CCC) by Phytoplant Research S.L. (Córdoba, ES). An Agilent liquid chromatography set-up (Model 1260, Pittsburgh, PA, United States) consisting of a binary pump, a vacuum degasser, a column oven, an autosampler and a diode array detector (DAD) equipped with a 150 mm length, 2.1 mm internal diameter, 2.7 mm pore size Poroshell 120 EC-C18 column was used for the quality control of the purified cannabinoids. The analysis was performed using water and acetonitrile both containing ammonium formate 50 mM as mobile phases. Flow-rate was 0.2 mL/min and the injection volume was 3 μL. Chromatographic peaks were recorded at 210 nm. All determinations were carried out at 35 °C. All samples were analyzed in duplicate. The result of Δ^9^-THCA-A purity 95.42% and THC impurity 1.32% was calculated as weight (%) versus a commercial standard from THC Pharm GmbH (Frankfurt, Germany) and Cerilliant (Round Rock, Texas, USA).

### Cell lines and luciferase assays

HEK-293T and 3T3-L1 cells cells were cultured in Dulbecco’s Modified Eagle’s Medium (DMEM) supplemented with 10% FBS, 2 mM L-glutamine and 1% (v/v) penicillin/streptomycin (Sigma-Aldrich, USA) and maintained at 37°C and 5% CO_2_ in a humidified atmosphere. HEK-293T (1 x 10^5^) cells were seeded in 24-well plates and transiently co-transfected with the indicated constructs (GAL4-PPARγ, GAL4-PPARδ, GAL4-PPARα and GAL4-luc) using Roti©-Fect (Carl Roth, Karlsruhe, Germany). After treatments, the luciferase activities were measured using Dual-Luciferase Assay (Promega, Madison, WI, USA).

### In vitro adipocyte and osteoblast differentiation

Human mesenchymal stem cells (MSCs) derived from bone marrow were seeded in α-MEM containing 10% FBS, 2 mM Glutamine, 1 ng/ml bFGF, and antibiotics, and adipocyte (AD) and osteoblast (OM) differentiation was performed as described (48). Treatment with RGZ (1 μM) and THCA-A (1, 5 and 10 μM) in the presence and the absence of T0070907 (5 μM) started at the same time as the differentiation process. After 7 or 14 days of differentiation, the mRNA was analyzed by qPCR and, after 21 days, adipogenesis and osteoblastogenesis were analyzed by Oil red O and Alizarin red staining respetively. The lipid accumulation and mineralization was quantified by removing the staining solution and absorbance was read at 540 and 405 nm, respectively.

### Animal studies

Six-week old male C57BL6 mice, obtained from Charles Rivers Laboratories (l’Arbresle, France), were pair-housed under constant conditions of light (12 hours of light/dark cycles) and temperature (22 ± 2 °C), with free access to food (see below) and water. All procedures concerning animal use were reviewed and approved by the Ethics Committee of the University of Cordoba and carried out in accordance with European Union Directive 2010/63/EU for the use and care of experimental animals.

At 8 weeks of age, mice were randomly assigned in two groups (N = 20) and fed either a high-fat diet (HFD), D12451 (Research Diets, New Brunswick, NJ, USA; 45%, 20%, and 35% calories from fat, protein and carbohydrate, respectively) or a standard diet (CD) (A04 SAFE Diets, Augy, France; 8,4%, 19,3% and 72,4% calories from fat, protein, and carbohydrate, respectively) during 15 weeks. Body weight (BW) gain, terminal BW, and daily energy intake were monitored once weekly along the first 12 weeks, and twice a week during treatment period. The latter was calculated from mean food ingestion per week using the energy density index provided by the manufacturer (3.34 kcal/g for CD or 4.73 kcal/g for HFD).

In order to assess the potential metabolic effects of Δ^9^-THCA-A, mice were treated daily by intraperitoneal injection of this compound (20 mg/Kg dissolved in ethanol/cremophor/saline 1:1:18) for three weeks, from week 12 onwards, in CD and HFD groups (n = 10/group). The amount of fat mass and adiposity in all set of experiments were measured by Quantitative magnetic resonance (QMR) scans, using the EchoMRI™ 700 analyzer (Houston, TX, USA, software v.2.0), before initiation of the treatments with Δ^9^-THCA-A and at the end of the experimental procedures. At this point, mice were euthanized and blood and brown adipose tissue (BAT), white adipose tissue (WAT) and liver were collected. Tissues were snap-frozen on dry ice and/or fixed in 4% formalin for further analysis of molecular expression and histology, respectively.

In another set of experiment, eight-week old male C57BL6 mice fed with CD were treated with Δ^9^-THCA-A (20 mg/Kg i.p.), with or without the selective PPARγ inhibitor, T0070907 (5 mg/Kg i.p.), for 3 weeks (n = 10/group). Pair-aged animals treated with vehicle served as control (n= 10/group).

### Immunohistochemistry and protein analysis by Western Blots

Liver and iWAT tissues were fixed in formalin for 24 hours, embedded in paraffin and sectioned. Liver tissue sections (5 μm) were stained with haematoxylin and eosin (H&E). Slides were evaluated for steatosis according to the Kleiner system. A semi-quantitative score was assigned to describe the extent of steatosis (0, <5%; 1, 5–33%; 2, 33–66%; and 3, >66%). IHC analysis of iWAT tissue sections (7 μm) was carried out as described previously (48). Briefly, antigen retrieval was performed in trypsin (pH 7.8) or 10 mM sodium citrate buffer (pH 6) and then incubated with F4/80 antibody (1:50; MCA497, Bio-Rad) or UCP-1 antibody (1:500; ab10983, Abcam, Cambridge, UK) overnight at 4 °C, respectively. Samples were analysed with a Leica DM2000 microscope and pictures were taken with a Leica MC190 camera. For Western blots, proteins were isolated from inguinal white (iWAT) adipose tissues and 30 μg of proteins were subjected to SDS/PAGE electrophoresis. Separated proteins were transferred (20V for 30 min) to polyvinyl difluoride membranes (PVDF) membranes that were probed probed with antibodies anti-UCP-1 (1:2000, Ab10983, Abcam) and α-tubulin (1:10.000; DM-1A, Sigma Aldrich). Differentiated 3T3-L1 cells in adipogenic medium were preincubated with either Δ^9^-THCA-A or RGZ and treated with 50ng/ml TNFα for 30min and the expression of PPARγ Ser272 and total PPARγ analysed with the antibodies anti-PPARγ Ser272 (1:200, bs-4888R, Bioss, Woburn, MA, USA) and anti-PPARγ (D1:1.000, 2435, Cell Signaling Technology, Danvers, MA, USA). Protein levels were normalized to β-actin (1:5000 dilution, A5060, Sigma Aldrich). Membranes were washed and incubated with the appropriate horseradish peroxidase-conjugated secondary antibody for 1 h at room temperature and detected by chemiluminescence system (GE Healthcare Europe GmbH, Freiburg, Germany).

### Intraperitoneal glucose and insulin tolerance tests, and triglyceride determinations

The animals were ip injected with a bolus of 2 g of glucose per kg BW, after a 5 h period of food deprivation, and blood glucose levels were determined at 0, 20, 60 and 120 min after injection. For ITT, the animals were subjected to ip injection of 1 U of insulin (Sigma Aldrich) per kg body weight, after a 5 h fasting. Blood glucose levels were measured at 0, 20, 60 and 120 minutes. All glucose concentrations were measured using a handheld glucometer (Accu-Check Advantage^®^; Roche Diagnostics, Rotkreuz, Switzerland). In addition, serum triglyceride levels were assayed, using a GPO-POD assay kit (Triglyceride Liquid kit 992320, Quimica Analitica Aplicada SA, Tarragona, Spain).

### Determination of hormonal, metabolic and inflammatory markers

Circulating adipokine levels of Leptin, Insulin, Glucagon, Glucagon-like peptide 1 (GLP-1) and Adiponectin were measured using quantitative Bio-Plex Pro™ Mouse Diabetes 8-Plex immunoassay (#171F7001M; Bio-Rad Laboratories, Hercules, CA, USA) and Bio-Plex Pro Mouse Diabetes Adiponectin assay #171F7002M (Bio-Rad Laboratories, Hercules, CA, USA) according to the manufacturer’s instructions. For proteome array, plasma samples from mice were pooled (n = 6 mice per group) and assayed for cytokine and adipokine expression. To study protein expression profiles, 100 μl plasma samples were used in the Proteome Profiler Mouse XL Cytokine Array and the Proteome Profiler Mouse Adipokine Array (R&D Systems, Minneapolis, MN, USA) according to the manufacturer’s protocols. Spot density was determined using Quick Spots image analysis software (R&D Systems).

### RNA-Seq and bioinformatic analysis

For each group, total RNA was extracted from iWAT, prepared, pooled, and run on an Agilent Bioanalyzer system to confirm quality (RNA integrity number >8). Transcriptome libraries were then constructed using poly-A selection with the TruSeq Stranded mRNA Library Prep Kit (Cat. No. RS-122-2101, Illumina, San Diego, CA, USA). In brief, 300 ng of total RNA from each sample was used to construct a cDNA library, followed by sequencing on the Illumina HiSeq 2500 with single end 50 bp reads and ~30 millions of reads per sample. The FASTQ files were pre-processed with Trimmomatic (v0.36) (52) to remove adapter sequences and aligned to the mm10 assembly of the mouse genome using HISAT2 (v2.1.0) (53). The counts per gene matrix were obtained from the alignments with featureCounts (v1.6.1) (54) using the in-built RefSeq annotation for the mm10 genome assembly. After filtering genes with less than 15 reads across samples, the raw counts were analyzed with DESeq2 (v1.20.0) (55) to obtain the regularized log transformed expression matrix and the differential expression analysis results. We used a threshold of an absolute fold change ≥ 2 and an adjusted P value ≤ 0.01 to consider a gene as differentially expressed in any comparison. Heatmaps were generated using the scaled mean of the regularized log expression for each group with the R package ComplexHeatmap (v1.20.0). The gene set enrichment analysis (GSEA) (56) and the over-representation analysis were performed using the R package ClusterProfiler (v3.10.1) (57). For GSEA, genes were pre-ranked using the log2 transformed fold change. The KEGG pathway database and Gene Ontology (Biological Process) annotation were used to group genes by biological function. All the P values were adjusted using the Benjamini and Hochberg correction to control the false discovery rate (FDR).

### Real-time PCR

Total RNA was isolated at day 7 or 14 of MSCs differentiation using the High Pure RNA Isolation kit (Roche Diagnostics). For tissues, total RNA was isolated using the Qiagen RNeasy Lipid Kit (Qiagen, Hilden, Germany). For real-time PCR analysis, RNA was reverse transcribed to cDNA using the iScript™ cDNA Synthesis Kit (Bio-Rad), and the cDNA generated was analysed by real-time PCR using the iQTM SYBR Green Supermix (Bio-Rad) using a CFX96 Real-Time PCR Detection System (Bio-Rad). Gene expression was normalized to HPRT or GADPH mRNA levels in each sample. The HPRT or GAPDH gene was used to standardize mRNA expression in each sample. The primers used are listed in Supplementary Table S2.

### Statistical analysis

In vitro data are expressed as mean ± SD with a minimal of 3 to 4 independent experiments. In vivo results are represented as mean ± SEM and the determinations were conducted with a minimal total number of 6-10 animals per group. Statistical analyses were performed on data distributed in a normal pattern, using Student’s t tests or ANOVA followed by Tukey’s test. P < 0.05 was taken as the minimum level of significance. Statistical analysis was performed using GraphPad Prism^®^ version 6.01.

## AUTHOR CONTRIBUTIONS

BP, RM and MEP conducted the *in vitro* experiments, the histopathology and the immuno-histochemistry studies. FRP coordinated and conducted, together with BP, IV, MASG, MEP and MJV, the whole set of *in vivo* experiments, as well as different of the analytical procedures related with those. MGR and MAC conducted the *in-silico* experiments and the bioinformatic analysis. BP, FRP, MAC and MGR were responsible for initial data analysis, figure preparation and statistical analysis. XN and CFV provided several batches of Δ^9^-THCA-A. MTS, GM and GA had a leading contribution in the design of *in vivo* studies, and an active role in the discussion and interpretation of the whole dataset. EM was responsible for the overall design of study, and major coordinator of the whole set of experiments. EM and MTS jointly wrote the manuscript as well as by the rest of the authors. All the authors take full responsibility for the work.

## ACKNOWLEDGEMENTS

This work was supported by grants SAF2017-87701-R (EM) and BFU2014-57581-P and BFU2017-83934-P (MTS) (Ministerio de Economía y Competitividad, Spain; co-funded with EU funds from FEDER Program); Project P12-FQM-01943 (M.T.-S.; Junta de Andalucía, Spain). CIBER Fisiopatología de la Obesidad y Nutrición is an initiative of Instituto de Salud Carlos III. None of the funding bodies played any role in the study design, data collection and analysis, the decision to publish, or the preparation of the manuscript. BP is a predoctoral fellow supported by the i-PFIS program, Instituto de Salud Carlos III (IFI15/00022; European Social Fund “investing in your future”).

