## Supplemental Data for "Tetrahydrocannabinolic Acid a (THCA-A) Reduces Adiposity and Prevents Metabolic Disease Caused by Diet-Induced Obesity"

[1]Instituto Maimónides de Investigación Biomédica de Córdoba (IMIBIC), Córdoba, Spain; [2]Departamento de Biología Celular, Fisiología e Inmunología, Universidad de Córdoba, Córdoba, Spain; [3]Hospital Universitario Reina Sofía, Córdoba, Spain; [4]Innovative Health Group, Madrid; [5]Emerald Health Biotechnology España, Córdoba, Spain: [6]Phytoplant Research, Córdoba, Spain; [7]Dipartimento di Scienze del Farmaco, Università del Piemonte Orientale, Novara, Italy; [8]Emerald Health Naturals, Vancouver, Canada.

### Methods

#### *Docking analysis*

Ligand docking, and binding properties were calculated by using the *AutoDock4 (1)* and the *Vina* software (2) with the virtual screening tool PyMOL (3). The receptor models used were the PDB references 5Y2O57 (4), 4EMA56 (5) and 5LGS58 (6). Search space for the docking was set around the binding sites described previously (7, 8).

#### *Time-resolved fluorescence resonance energy transfer (TR-FRET)*

Time-resolved fluorescence resonance energy transfer (TR-FRET) coactivator assay for PPARγ was performed by employing LanthaScreen™ kit (ThermoFisher Scientific®, #A15126) according to the manufacturer's protocol. Serial concentrations of THCA or rosiglitazone (RGZ) were incubated with GST-fused human PPARγ-LBD, terbium-labeled anti-GST antibody, and a fluorescently labeled PPARγ ligand peptides for 4 h in the dark at room temperature. FRET signal was measured by excitation at 340 nm and emission at 520 nm for fluorescein and 495 nm for terbium using a FlextStation3 Benchtop Multi-Mode microplate reader in 384-well plate format. Ligand peptides for PPARγ containing a N-terminal FITC label were TRAP-220 (#PV4549) and PGC1α (#PV4421) as coactivators and NCoR (#PV4624) and SMRT (#PV4423) as corepressors and they were obtained from ThermoFisher Scientific®.

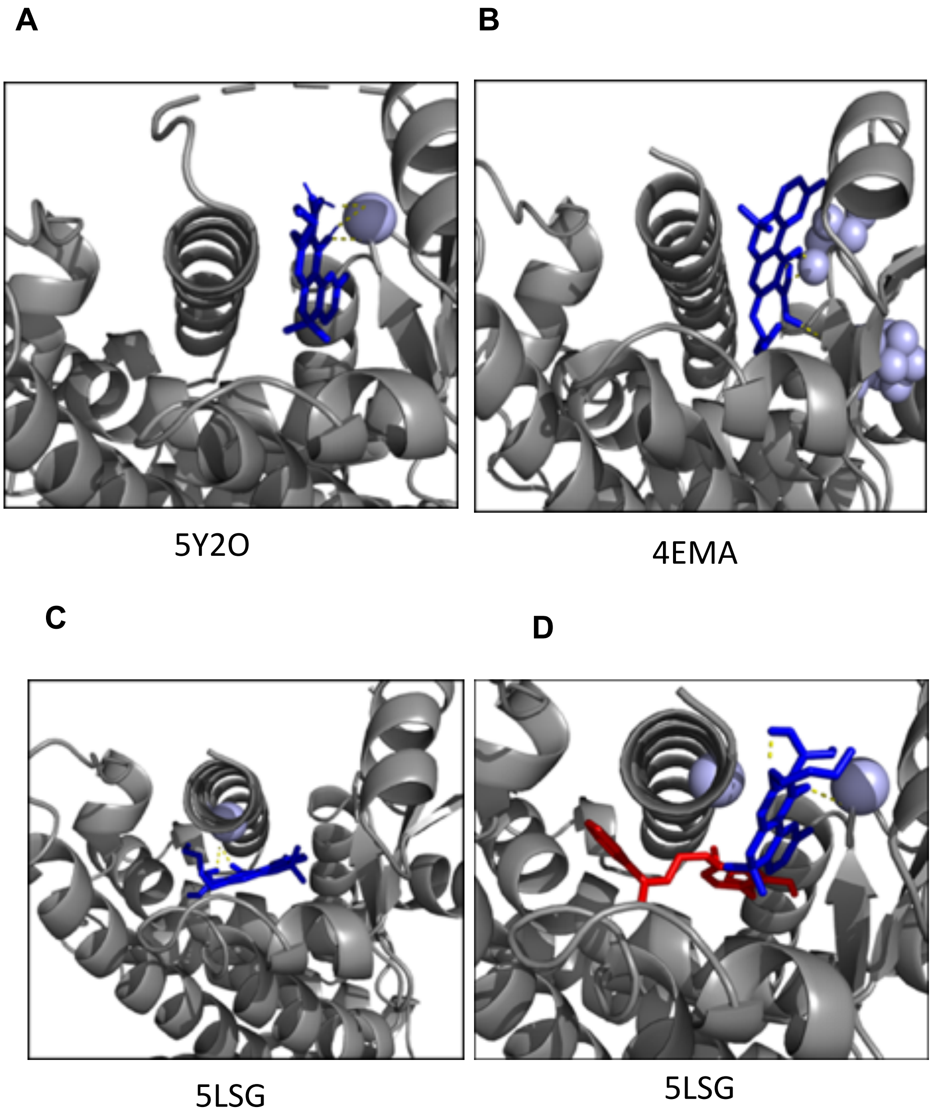


**Supplementary Figure S1.** (**A**) PPARγ LBD structure 5Y2O ser342 bound to $\Delta^9$-THCA-A (blue) B.E. AutoDock Kcal/mol= -8.8; B.E. VINA Kcal/mol= -7.4; Predicted Ki 336,79 nM. (**B**) PPARγ LBD structure 4EMA ser342 and L340 bound to $\Delta^9$-THCA-A (blue) B.E. AutoDock Kcal/mol= -7.45; B.E. VINA Kcal/mol= -7.4; Predicted Ki 3,49 μM. (**C**) PPARγ LBD structure 5LSG ser289 in Helix 3 bound to $\Delta^9$-THCA-A in the orthosteric site (blue), B.E. AutoDock Kcal/mol= -8.95; B.E. VINA Kcal/mol= -7.9; Predicted Ki 273,22 nM. (**D**) PPARγ LBD structure 5LSG bound to RGZ in the orthosteric site (red), and $\Delta^9$-THCA-A in the alternative site (blue).

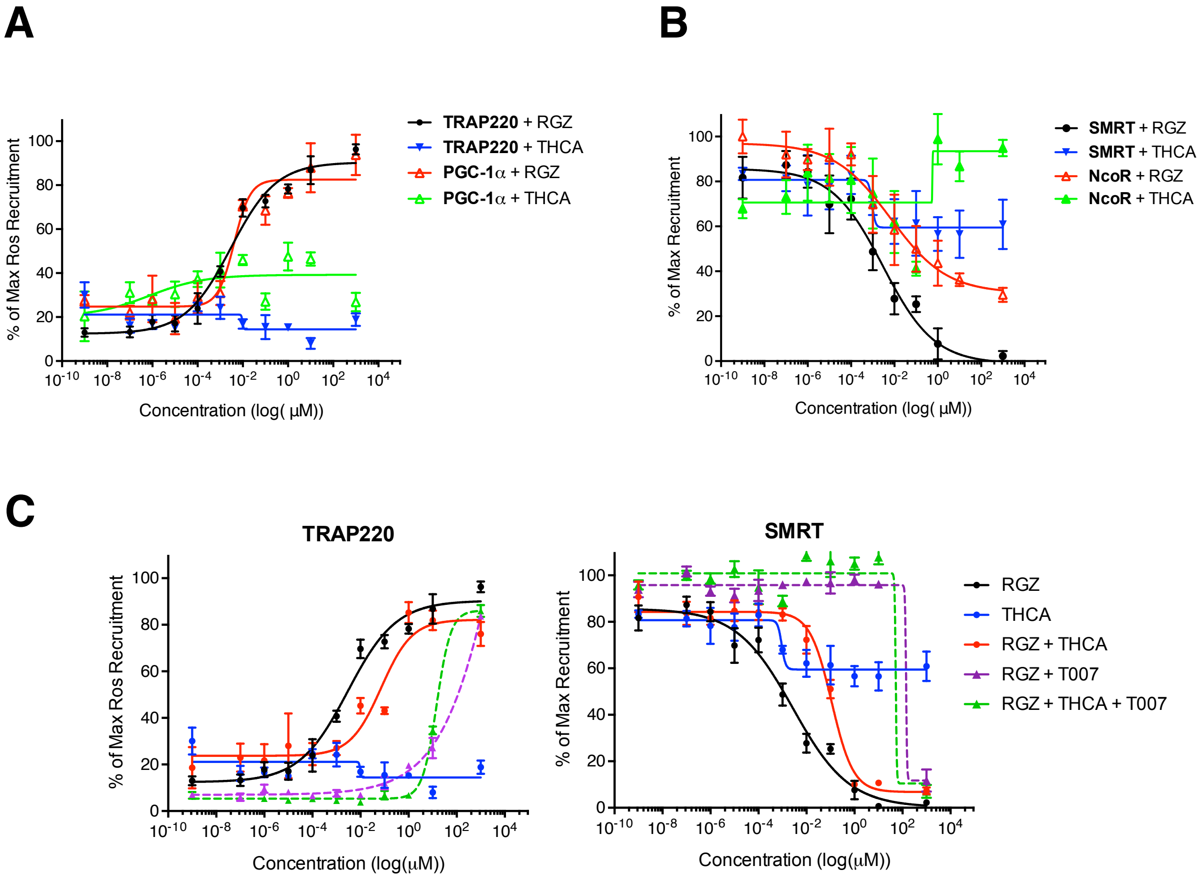


**Supplementary Figure S2. Effects on PPARγ-ligand interactions in presence of Δ$^9$-THCA-A in LanthaScreen assays. (A)** TR-FRET assay was employed to study coactivator peptide recruitment to the human PPARγ LBD in response to RGZ or Δ$^9$-THCA-A. Data are expressed as a percentage of the maximum RGZ response. Three experiments were performed, and representative graphs are shown. **(B)** Corepressor peptide displacement to human PPARγ LBD in response to RGZ or Δ$^9$-THCA-A was examined by TR-FRET assay. Data are expressed as a percentage of the maximum recruitment in the absence of ligand. Representative data from three experiments are shown. **(C)** Alternative site binding for Δ$^9$-THCA-A in PPARγ was tested by using increasing concentrations of RGZ or Δ$^9$-THCA-A both in the absence or the presence of 5µM T0070907 PPARγ antagonist. TRAP220 data are expressed as a percentage of the maximum RGZ response, while SMRT data are expressed as a percentage of the maximum recruitment in the absence of ligand. Representative graphs from three experiments are shown.

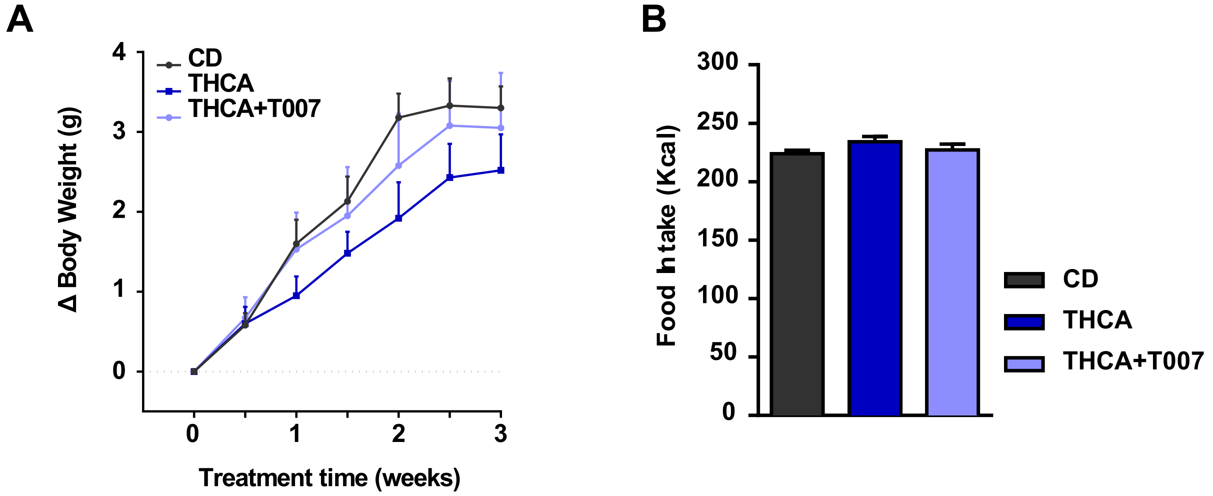


95 **Supplementary Figure S3.** Effect of 3 weeks treatment of $\Delta^9$-THCA-A combined or not with of T0070907 on body weight gain and food intake. **(A)** BW gain, and **(B)** Total calorie intake (Kcal) during the treatment, for the three experimental groups.

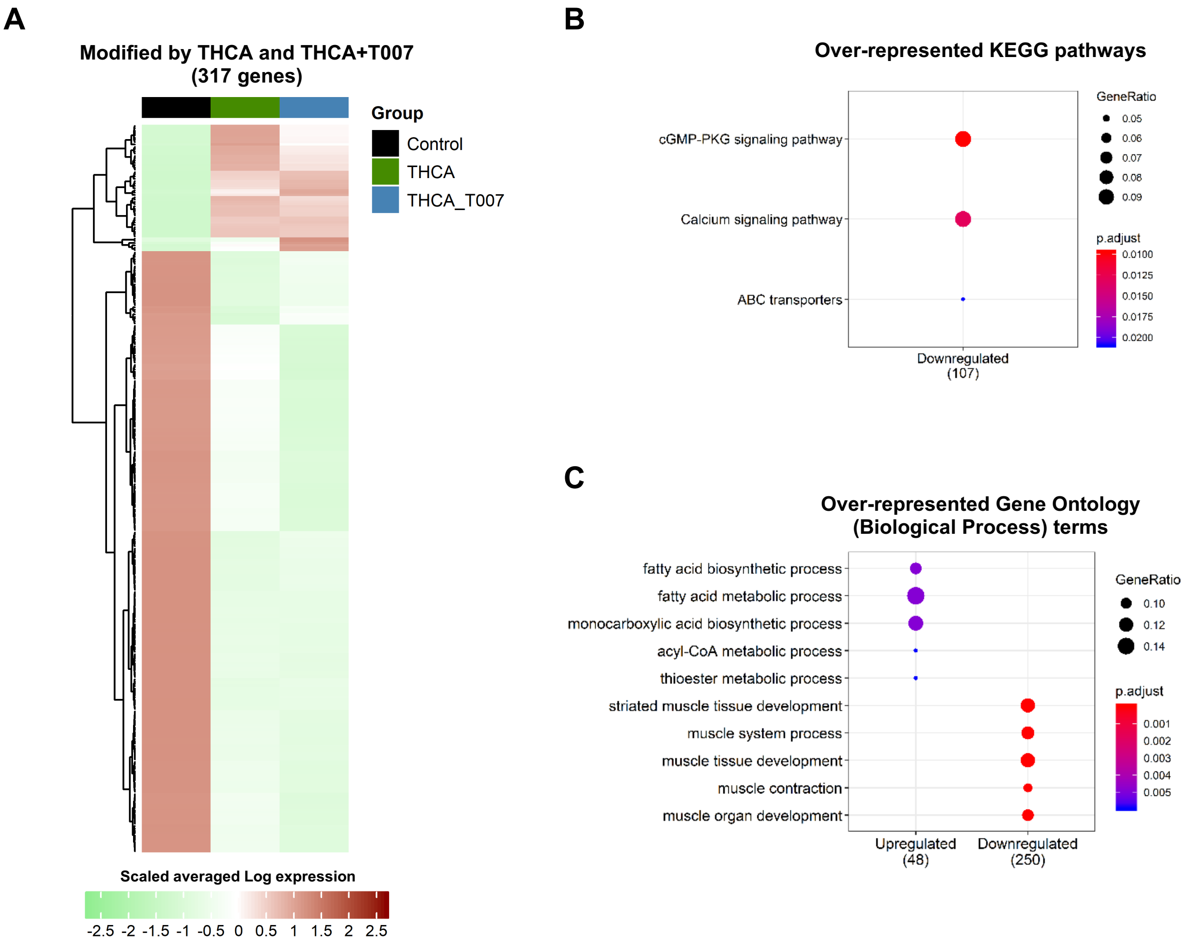


**Supplementary Figure S4.** (A) Heatmap of the 317 genes that are modified in the same direction by both $\Delta^9$-THCA-A and $\Delta^9$-THCA-A+T0070907 treatments. The color represents the scaled mean of log transformed expression. The column annotations indicate the sample group. (B) Over-represented KEGG pathways and Gene Ontology (Biological Process) terms in the clusters of up or down regulated genes by both treatments. The presence of a point indicates the over representation (Fisher Exact Test Adjusted P $\leq$ 0.05) of a pathway or term (Y axis) in a group of genes (X axis).

**Supplementary Table 1: Primers used in this study:**

| **Gene** | **Forward (5´ → 3´)** | **Reverse (5´ → 3´)** |
|---|---|---|
| **h-PPARγ2** | GCGATTCCTTCACTGATACACTG | GAGTGGGAGTGGTCTTCCATTAC |
| **h-LPL** | GGCGCTACCTTGAGATAGAGTTCTG | TGTTTTCTACAGGGTGCTTTAGATGAC |
| **h-aP2a** | CCAGGAATTTGACGAAGT | TCTCTTTATGGTGGTTGATT |
| **h-CEBPA** | CCTTGTGCCTTGGAAATGCAAAC | CTGCTCCCCTCCTTCTCTCA |
| **h-ADIPOQ** | CATGACCAGGAAACCACGACTC | CCGATGTCTCCCTTAGGACCA |
| **h-RUNX2** | TGGTTAATCTCCGCAGGTCAC | ACTGTGCTGAAGAGGCTGTTG |
| **h-ALP** | CCAACGTGGCTAAGAATGTCATC | TGGGCATTGGTGTTGTACGTC |
| **h-SP7** | AGCCAGAAGCTGTGAAACCTC | AGCTGCAAGCTCTCCATAACC |
| **h-IBSP** | AGGGCAGTAGTGACTCATCCG | CGTCCTCTCCATAGCCCAGTGTTG |
| **h-HPRT** | ATGGGAGGCCATCACATTGT | ATGTAATCCAGCAGGTCAGCAA |
| **m-TNFα** | CTACTCCCAGGTTCTCTTCAA | GCAGAGAGGAGGTTGACTTTC |
| **m-ICAM1** | GTGGCGGGAAAGTTCCTG | CGTCTTGCAGGTCATCTTAGGAG |
| **m-CD4** | TCCTTCCCACTCAACTTTGC | AAGCGAGACCTGGGGTATCT |
| **m-CXCL16** | CGTTGTCCATTCTTTATCAGGTTCC | TTGCGCTCAAAGCAGTCCA |
| **m-CCL22** | AAGACAGTATCTGCTGCCAGG | GATCGGCACAGATATCTCGG |
| **m-CXCR5** | ACTCCTTACCACAGTGCACCTT | GGAAACGGGAGGTGAACCA |
| **m-CXCR2** | CACCGATGTCTACCTGCTGA | CACAGGGTTGAGCCAAAA |
| **m-GAPDH** | TGGCAAAGTGGAGATTGTTGCC | AAGATGGTGATGGGCTTCCCG |
